# Methadone alters transcriptional programs associated with synapse formation in human cortical organoids

**DOI:** 10.1101/2022.11.04.515240

**Authors:** Ila Dwivedi, Dan Zhou, Andrew B. Caldwell, Shankar Subramaniam, Gabriel G. Haddad

**Affiliations:** Department of Pediatrics, School of Medicine, University of California, San Diego, La Jolla, CA, USA; Department of Neurosciences, School of Medicine, University of California, San Diego, La Jolla, CA, USA; Department of Cellular and Molecular Medicine, School of Medicine, University of California, San Diego, La Jolla, CA, USA; Department of Bioengineering, University of California, San Diego, La Jolla, CA, USA; Department of Nanoengineering, University of California, San Diego, La Jolla, CA, USA; Department of Computer Science and Engineering, University of California, San Diego, La Jolla, CA, USA; Rady Children’s Hospital, San Diego, CA, USA

**Keywords:** Opioid Use Disorder, Methadone, Synapses, Brain Development, Organoids, Transcription

## Abstract

Opioid use disorder (OUD) among pregnant women has become an epidemic in the United States. Pharmacological interventions for OUD involve methadone, a synthetic opioid analgesic that attenuates withdrawal symptoms and behaviors linked with maternal drug abuse. However, methadone’s ability to readily accumulate in neural tissue, and cause long-term neurocognitive sequelae, has led to concerns regarding its effect on prenatal brain development. We took advantage of human cortical organoid (hCO) technology to probe how this drug impacts the earliest mechanisms giving rise to the cerebral cortex. To this end, we conducted bulk mRNA sequencing of 2-month-old hCOs derived from two cell lines that were chronically treated with a clinically relevant dose of 1μM methadone for 50 days. Differential expression and gene ontology analyses revealed a robust transcriptional response to methadone associated with functional components of the synapse, the underlying extracellular matrix (ECM), and cilia. Further unsupervised co-expression network and predictive protein-protein interaction analyses demonstrated that these changes occurred in concert, centered around a regulatory axis consisting of growth factors, developmental signaling pathways, and matricellular proteins. Our results demonstrate that exposure to methadone during early cortico-genesis fundamentally alters transcriptional programs associated with synapse formation, and that these changes arise by modulating extra-synaptic molecular mechanisms in the ECM and cilia. These findings provide novel insight into methadone’s putative effect on cognitive and behavioral development and a basis for improving interventions for maternal opioid addiction.

## INTRODUCTION

In the past two decades, Opioid Use Disorder (OUD) among pregnant women has become an epidemic in the United States (1–6). The rapid rise in rates of maternal opioid related diagnoses has been accompanied by a parallel surge in the percentage of expectant mothers seeking treatment for addiction, dependence, and abuse (2,5,7). The standard of care for maternal OUD is Medication-Assisted Treatment (MAT) with methadone (2,3,5), a synthetic mu (μ)-opioid analgesic that minimizes deleterious opioid withdrawal symptoms and risk-taking behaviors that lead to relapse or overdose (8–10).

Despite its utility in adults, methadone’s ability to readily enter fetal circulation and accumulate in neural tissue has led to concerns regarding its effects on fetal brain development (8,11). Clinically, prenatal methadone exposure is linked with the increased incidence and severity of Neonatal Abstinence Syndrome (NAS), which is characterized by central nervous system hyperirritability and autonomic nervous system dysfunction (5,8,12–14). Exposed infants also exhibit long-term psychomotor and cognitive sequelae such as impaired learning, memory, and motor skills, as well as depression and anxiety (15–20).

Aiming to uncover the underlying cause of these impairments, diffusion tensor imaging studies of neonates and adults prenatally exposed to methadone uncovered microstructural changes in cerebral white matter tracts (20–23). Early studies of methadone also showed that exposure diminishes neurotransmitter content, uptake, and release (24,25) as well as synaptogenesis (26,27). Treatments of rat neuronal cultures with other μ-opioid receptor agonists like morphine yielded reduced neurite outgrowth and pre- and post-synaptic puncta densities (28). Morphine, endogenous μ-opioids, and methadone were also shown to affect central and peripheral neuronal excitability and communication (29–32).

Nevertheless, most studies assessing methadone’s effects on fetal neurodevelopment have been limited by duration of exposure, model organism, or subject age (33). Investigations have largely been conducted in murine models, whose developmental timelines differ significantly from humans (34–37). The few studies done in human subjects have been postnatal and complicated by opioid-based pharmacotherapies for NAS, length of MAT, and maternal pathophysiology. Moreover, human fetal tissues are in short supply due to ethical and logistical complications. Consequently, methadone’s effects on human fetal brain development have remained largely uncharted.

To address these limitations, we utilized human iPSC-derived three-dimensional models of cortical development called cortical organoids (hCOs) (38,39). These contain multiple cell types and undergo spatial organization characteristic of the *in vivo* fetal cortex, eliminating postnatal factors that have confounded previous studies of prenatal methadone exposure (39–47). As prior studies of methadone and other μ-opioids had been linked to deficits in neural connectivity and communication, we hypothesized that exposure to the drug during cortical development would alter fundamental mechanisms underlying synaptogenesis in these hCOs. To study these potential changes, we conducted bulk mRNA sequencing of 2-month-old hCOs chronically treated with a therapeutically relevant concentration of 1μM methadone (8,48–50) for 50 days, a period leading up to the onset of synapse formation (38,39). This methodology modeled a clinical scenario of MAT beginning in the first trimester of pregnancy and enabled us to dissect how methadone alters molecular mechanisms underlying neural development in the fetal cortex.

## METHODS AND MATERIALS

### Human cortical organoid generation and treatment with methadone

hCOs were generated and maintained using a protocol previously described by Muotri and colleagues (Supplementary Methods) (38,39). iPSC lines (A & B) derived from two healthy individuals were used to generate the cortical organoids.

Methadone (Sigma-Aldrich, St. Louis, MO) was dissolved in sterile nuclease-free water (Invitrogen, Waltham, MA) and diluted to a working concentration of 1 μM in fresh stage-specific medium each day of media change. Treatment began on Day 9 of organoid culture, the first day of the neural proliferation stage, up to Day 60. Nuclease-free water was used as a vehicular control.

### Cortical organoid collection and RNA isolation

Cortical organoids were collected 2 months (60 days) after beginning organoid culture. Each well of hCOs was a separate biological replicate for a given treatment condition (i.e., treated versus untreated). hCOs in each well were aspirated in 1 mL of medium and centrifuged at 3000g for 5 minutes at room temperature. The pellet was resuspended in 1 mL cold (4°C) DPBS (Corning, Corning, NY) and centrifuged again at 4°C and 21,000g for 10 minutes. Pellets were snap frozen in 1.5 mL Eppendorf tubes using dry ice and stored at −80°C prior to RNA extraction. RNA was extracted from frozen organoid pellets using the Direct-Zol Miniprep Plus Kit (Zymo, Irvine, CA) according to the manufacturer’s instructions.

### RNA-sequencing data generation

Total RNA was sent to the UC San Diego Institute for Genomic Medicine (IGM) for quality assessment, library preparation, and sequencing. Only samples with RNA Integrity Numbers (RIN) > 7 were selected for library preparation. PolyA+ selected libraries were prepared with the TruSeq mRNA Stranded Library Prep Kit with TruSeq UDI96 indexed adaptors (Illumina, San Diego, CA). Samples were multiplexed and sequenced on the Illumina NovaSeq 6000 S4 to produce approximately 100 million, 100 base pair, paired-end reads per sample. Three control and three methadone-treated samples were sequenced from cell line A, and four control and four treated samples from cell line B. RINs and total sequenced reads per sample are given in Table S1.

### RNA-sequencing differential expression analysis

Raw fastq file quality assessment and read alignment to the hg19 genome (GRCh37, RefSeq GCF_000001405.13) (51) were performed through the *FastQC* (v1.0.0) (52) and RNA-Seq Alignment (*STAR*, v2.0.2) (53) applications, respectively, in the Illumina BaseSpace Sequence Hub. Mapped reads were assigned to genomic meta-features (genes) using the Rsubread (v2.6.4) (54) package *featureCounts* in R. Expression level filtering was performed using the *edgeR* (v3.34.1) (55) function filterbyExpr. TMM normalization factors (56) were also calculated using *edgeR*. Differential expression analysis was conducted using the R package *limma* (v3.48.3) (57,58) by incorporating cell line and treatment condition as covariates in the linear model and contrasting methadone treated versus untreated control samples. Genes were ranked in order of evidence for differential expression using the empirical Bayes (eBayes, t-value) method (n = 17,651). Significantly differentially expressed genes (DEGs) were selected based on the confident effect size of their log_2_(Fold Change) values at FDR < 0.05. This was represented by a “confect score” calculated by *TopConfects* (v1.8.0) (59) in R. Genes with |Confect Score| ≥ log_2_(1.5) were considered DEGs.

### Gene ontology and gene set enrichment analysis

To identify transcriptional signatures associated with specific cellular components, gene set enrichment analysis (GSEA) (60) was performed using the *fGSEA* (v1.18.0) (61) package in R with the Gene Ontology Cellular Component (GO-CC) database (62,63) as a reference and all eBayes ranked genes as input. The top 20 terms with the lowest FDR-adjusted p-values were selected and sorted by their normalized enrichment scores (NES).

To determine which molecular functions the DEGs associated with each top cellular component were enriched for, we applied a hypergeometric test using the GO-Molecular Function (GO-MF) database in the *GOrilla* web application (64) using eBayes ranked genes as background (p < 10^−3^). Resulting GO-MF terms were arranged hierarchically and non-redundantly by semantic similarity via the *REVIGO* web-application (65). *REVIGO* significance values and term hierarchy were used to visualize GO-MF enrichment as a circle plot using the *CirGO* package (66) in *Python* (Data File S1).

The molecular role of each synaptic DEG was determined based on its association with enriched GO Molecular Function categories and through manual searches using the *OMIM* (67,68), *GeneCards* (69,70), and *NCBI Gene* (71) databases. DEGs were grouped into categories based on the representative terms found in the *CirGO* output. ECM-associated DEGs were categorized by their overlap with Core- or Matrisome-Associated genes in *MatrisomeDB* (72). Types of genes in each category were then grouped using the descriptions in reviews written Naba and colleagues (72–75), who generated the *MatrisomeDB*. Major collagen (76) and proteoglycan (77,78) types were identified based on descriptions in the literature. As before, *OMIM, GeneCards*, and *NCBI Gene* databases helped categorize genes not identified in these sources (Data File S2).

### Modular gene co-expression and protein-protein interaction network analysis

Co-expression modules were identified using the Bioconductor *cemitool* package (79) in R, using eBayes ranked genes as input. Their counts were corrected for cell line differences using the removebatcheffects function and transformed using the voomwithqualityweights function in *limma*. GO-CC *fGSEA* was performed on each module, and DEGs from the top two modules with the greatest similarity in cellular component enrichment were combined for network analysis.

Edges between DEGs from the first two modules were acquired from *STRINGDB* (v.11.0) (80) and filtered to include only physical interactions (medium threshold = 0.7). Edges and nodes for each network were imported into *Cytoscape* (81), and top hub genes ranked by EcCentricity score (82) were identified using the *CytoHubba* application (83). *MCODE* (84) identified highly interconnected clusters of proteins in each network using default network scoring and cluster finding parameters.

### Upstream Regulator Analysis

Ingenuity Pathway Analysis (*IPA*; Qiagen Inc., Hilden, Germany) (85) was used to identify upstream regulators of synapse, cilia, and ECM associated DEGs from the first two co-expression modules. Each regulator was assigned a p-value representing the overlap between DEGs and known targets, and a z-score to infer regulator activation states. Endogenous molecules (excluding exogenous toxicants, drugs, and reagents) were ranked in order of significance (Data File S3).

## RESULTS

### Chronic methadone induces a robust transcriptional response in 2-month-old hCOs

Two-month-old hCOs generated from two iPSC cell lines, A and B, exhibited robust transcriptional responses to 50 days of chronic treatment with 1μM methadone (Figure 1A). 4165 DEGs were detected in line A, while 1018 DEGs were identified in line B (|Confect| ≥ Log_2_(1.5), FDR < 0.05) (Figure S1A). For each cell line, control samples were differentiated from methadone treated samples by unsupervised clustering according to their expression profiles (Figure S1B).

**Figure 1.**
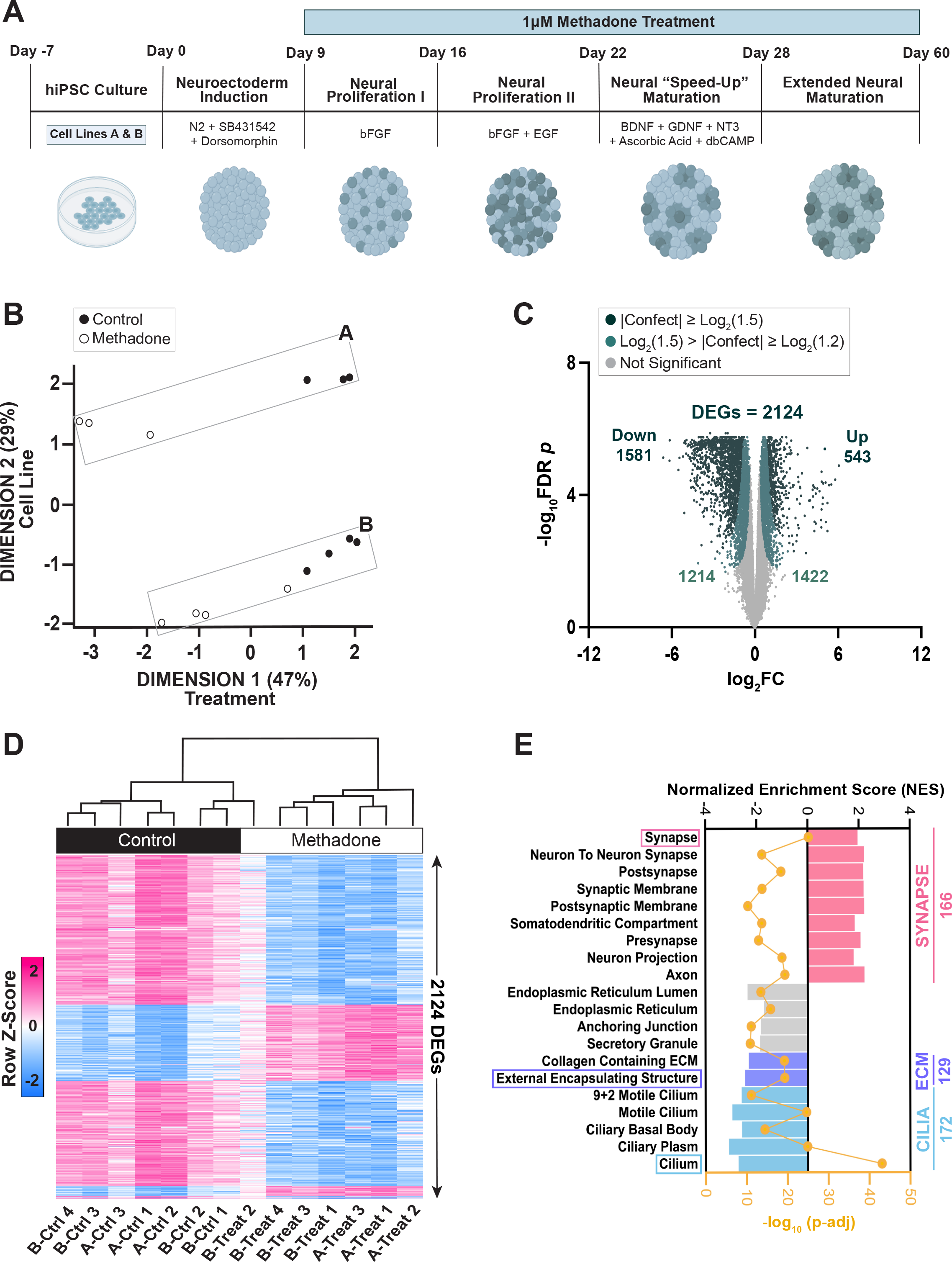
Methadone elicits a robust transcriptional response in 2-month-old hCOs. **(A)** Timeline of cortical organoid generation and methadone treatment. **(B)** Multi-dimensional scaling of TMM-normalized expression data and metadata from control and methadone-treated hCO samples derived from cell lines A & B. **(C)** RNA-seq volcano plot of all (eBayes ranked) genes, distinguished by their confident effect size ‘confect’ score cutoffs. Genes with |Confect| ≥ log_2_(1.5) and FDR < 0.05 were considered DEGs (DEGs = 2124), after adjusting for cell-line differences. For each gene, log_2_ (Fold Change) effect size values are shown on the x-axis and Benjamini-Hochberg adjusted absolute log_10_ p-values are along the y-axis. **(D)** Heat map depicting the sample-level expression (z-scores) of the 2124 DEGs across both cell lines. **(E)** Top 20 fgsea enriched GO Cellular Component gene sets based on absolute log_10_ FDR-adjusted p-values, ranked according to the direction (positive or negative) of their normalized enrichment scores (NES) and grouped by cellular component. Values to the right of the graph indicate the number of DEGs associated with each of the top 3 enriched GO-CC gene sets.

Despite differences in the magnitude of response to methadone between the cell lines, multi-dimensional scaling revealed that methadone treatment was the primary source of sample-level variation (Figure 1B). Furthermore, 76.3% of the DEGs in cell line B overlapped with those in cell lines A, demonstrating a high degree of similarity in response to methadone treatment (Figure S1C). This information enabled us to incorporate cell line as a covariate into the differential expression linear model and account for baseline transcriptional differences between individuals.

This method yielded 2124 DEGs whose expression was altered due to methadone alone (Figure 1C). Almost all samples from both cell lines clustered together according to treatment condition based on their expression of these DEGs (Figure 1D). Although B-Treat 2 clustered with control samples, its high quality and read counts precluded its exclusion from further analyses (Figure 1D; Table S1). Furthermore, there was significant overlap between the final 2124 DEGs and those from each individual cell line. Approximately 72% of the final 2124 DEGs intersected with those in line A, while 93% of the DEGs in line B overlapped with the final DEGs, indicating that the transcriptional response to methadone was largely preserved even after adjusting for cell line etiology (Figure S1D).

### Methadone alters the expression of pre- and post-synaptic functional components

Since we had hypothesized that methadone would affect synapse formation, we began by determining how the observed transcriptional response was associated with the cellular anatomy of our organoid system. Pre-ranked gene set enrichment analysis (GSEA) using the Gene Ontology Cellular Component (GO-CC) database revealed the significant enrichment of gene sets associated with neuronal synapses (Figure 1E). We noted sets linked to both the pre- and post-synapse, distinguished by terms such as “Axon” or “Somatodendritic Compartment”, respectively. All sets had positive normalized enrichment scores (NES), trending towards transcriptional upregulation in the synapse following methadone exposure.

To identify which aspects of synaptic biology were specifically affected by methadone, we performed a ranked GO-Molecular Function (GO-MF) enrichment analysis of the synaptic DEGs and summarized the resulting non-redundant ontology terms hierarchically (Figure 2A; Data File S1). We observed alterations in all aspects of synaptic biology, including the pre-synaptic trafficking, release and synthesis of hormones and neurotransmitters, the postsynaptic reception and response to these signaling molecules, and intracellular cytoskeletal scaffolding (86–88). There were 166-synapse associated DEGs, of which 58 were identified as pre-synaptic, 64 as post-synaptic, and 44 as linked to both terminals (Figure 2B; Data File S2). To further resolve the identities of these genes, we categorized them functionally according to the GO-MF, *OMIM, GeneCards*, and *NCBI Gene* databases. Most pre-synaptic DEGs were involved with vesicular trafficking, while post-synaptic DEGs were primarily receptors or signal transduction molecules. The 44 DEGs associated with both terminals were primarily structural, involved with cytoskeletal integrity, cell-cell adhesion, or extracellular matrix (ECM) composition.

**Figure 2.**
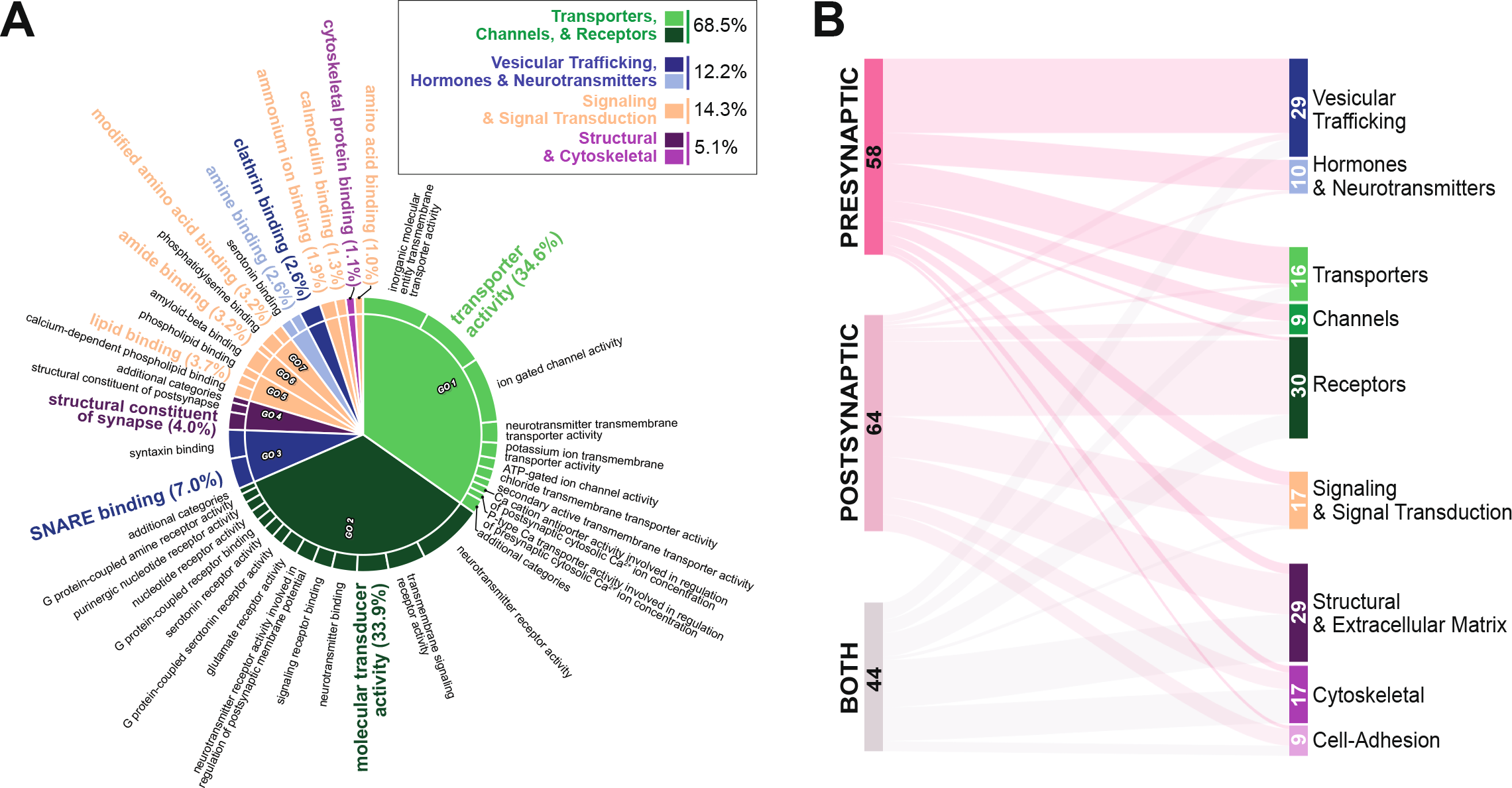
Methadone alters the expression of pre- and post-synaptic functional constituents. **(A)** Molecular function ontology of synapse-specific DEGs compared, determined via a hypergeometric test using a background of eBayes-ranked expressed genes and a GO gene set significance threshold of p < 10^−3^. “Parent” or representative gene sets (molecular function categories) are presented in the inner circle and emphasized with large, colored font. Semantically similar descendants of these parents are represented in black font. Terms in each category are sorted in descending order by absolute log_10_ p-values. Percentages indicate the relative membership of each parent gene set based on the number and significance of the constituent descendant terms. Gene sets were broadly summarized and colored according to the categories shown in the boxed insert. **(B)** Molecular functions of 166 synapse associated DEGs. Node values and sizes, as well as edge thicknesses represent the number of DEGs belonging to each GO cellular component category (pre-synapse, post-synapse, both) or molecular function.

### Methadone induces transcriptional changes in an ECM regulatory hierarchy

Consistent with the identities of the 44 shared pre- and post-synaptic DEGs, we detected two GO-CC gene sets with negative NES values associated with the ECM (“external encapsulating structure” and “collagen containing ECM”) (Figure 1E). The ECM is a critical component of the tetrapartite synapse, which otherwise consists of the presynaptic bouton, postsynaptic terminal, and supporting glial cells (89–91). Given the ECM’s role in synapse formation and maintenance, we studied how its composition and function had been affected by methadone treatment. To this end, we conducted a ranked GO-MF enrichment analysis of the 129 ECM-associated DEGs and summarized the resulting non-redundant terms hierarchically. Four distinct functional categories of matrix structural, binding, catalytic, or signaling molecules were enriched, encompassing the ECM regulatory hierarchy (Figure 3A; Data File S1) (92–94).

**Figure 3.**
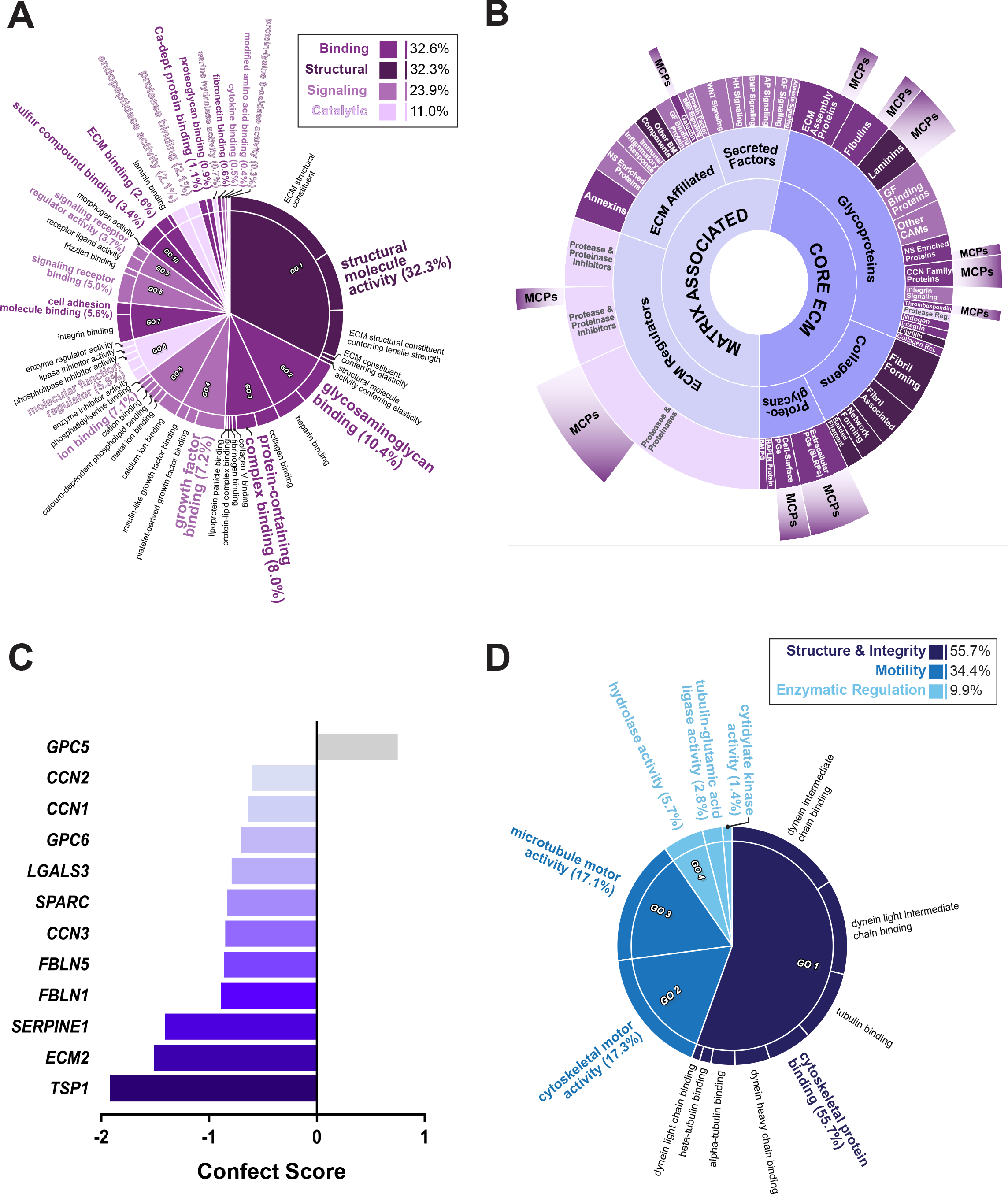
Methadone induces transcriptional changes associated with ECM and ciliary structure and function. **(A)** Enriched molecular function ontology of ECM-specific DEGs compared to a background of eBayes-ranked expressed genes. Parent and descendant terms were determined and visualized as described in Figure 2A. Broad categories of enriched ECM gene sets are summarized in the boxed insert. **(B)** Categorization of ECM-associated DEGs. The inner two circles group molecules according to their designations in *MatrisomeDB*. The third circle organizes molecules in each Matrisome category based on their functions. The outermost bars indicate the number of matri-cellular proteins (MCPs) in each category. Segment sizes correspond to the number of DEGs in each category. **(C)** All MCPs classically identified in the brain that were differentially expressed in response to methadone. Genes are ranked by their confect score values. **(D)** Enriched molecular function ontology of ciliary-DEGs compared to a background of eBayes-ranked expressed genes. ECM, extracellular matrix; CAM, cell adhesion molecule; GF, growth factor; NS, nervous system, Reg, regulation; Rel, related; HAPLN, hyaluronan and proteoglycan link; BM, basement membrane; HH, hedgehog; AP, angiopoetin.

To parse these DEGs further, we sorted them according to *MatrisomeDB*, a collection of genes defined as core (collagens, proteoglycans, and other glycoproteins) or associated (ECM regulators, secreted factors, or other affiliated molecules) matrix proteins (72–75). Methadone treatment disrupted the expression of genes belonging to each of these designations (Figure 3B; Table S2; Data File S2). Glycoproteins (other than proteoglycans) made up over half of the core matrix proteins in our dataset (55.6%, n = 35/63). These included structural molecules like laminins, cell adhesion molecules, and non-structural matrix modulators belonging to the CCN or thrombospondin family of proteins (92–94). ECM regulators also constituted almost half (n = 31/66, 47%) of the matrix-associated (non-core) proteins. These were primarily matrix proteases and their inhibitors (n = 23/31, 74%), including a family of matrix metalloproteinases (MMPs or ADAMTS) that modulate ECM composition by hydrolyzing its components (95,96).

Notably, over a quarter of all ECM DEGs encoded known and proposed matricellular proteins (MCPs) (n = 37/129, 28.7%) (Figure 3B; Data File S2) (97–105). MCPs are non-structural proteins that modulate ECM composition and integrity through interactions with structural proteins, proteases, cell surface receptors, and growth factors (97,99,101). In the developing brain, MCPs regulate mechanisms of cellular maturation, proliferation, migration, axonal guidance, and synapse formation. Among the MCP families classically found in the brain, thrombospondins (thrombospondin1, *TSP1*), *SPARC* (including *Hevin/SC1/ECM2*), the Cellular Communication Network Factors (*CCN1-3* or *CYR61*, *CTGF*, and *NOV*), glypicans (*GPC-5* and *-6*), galectins (*LGALS3*), plasminogen activator inhibitor (*SERPINE1/PAI-1*), and fibulins (*FBLN1* and *5*) were all differentially expressed due to methadone (Figure 3C; Data File S2) (106–110). Of these, *TSP1* exhibited the greatest magnitude of change in response to treatment (Figure 3C; confect score = −1.92, FDR < 0.05). Altogether, these changes indicated that methadone alters crucial components of the ECM regulatory hierarchy that are necessary for developmental synapse formation.

### Methadone disrupts the expression of genes involved in ciliary integrity and motility

Unexpectedly, changes to the ECM were also accompanied by the enrichment of cilium-associated gene sets (Figure 1E). Like the ECM, the 172 ciliary DEGs were mostly downregulated, indicated by the negative NES values. These DEGs could be separated into three groups encoding motor proteins (e.g., kinesins or dyneins), cytoskeletal proteins contributing to projection integrity, and enzymes potentially regulating cytoskeletal or motor protein polymerization (Figure 3D; Data File S1) (111–113). This data reveals a concurrent, directional response to methadone in transcriptional programs informing ECM and ciliary structure and function. This pattern of synaptic, ECM, and ciliary enrichment was observed in cell lines A and B individually as well, reinforcing a consistent and parallel change in synaptic and extra-synaptic biology in response to drug treatment (Figure S2A-C).

### Synaptic, ECM, and ciliary DEGs are highly co-expressed and encode proteins that physically interact

We next investigated the relationship between DEGs belonging to the synaptic, ciliary, and ECM compartments. Comprehensive modular co-expression analysis of eBayes ranked genes yielded 14 modules of highly co-expressed genes. Of these, the top two modules retained the greatest DEG membership ratios (module M1 = 1572/6380 and module M2 = 330/4382) (Figure 4A). GO-CC GSEA revealed that M1 and M2 were significantly enriched for gene sets associated with the synapse, ECM, and cilia (Figure 4B, Table S3). This ontological overlap between M1 and M2 allowed us to use the intersection of their DEGs for further analyses, and indicated that changes to the synapse were not occurring in isolation, but in concert with changes to the ECM and cilia.

**Figure 4.**
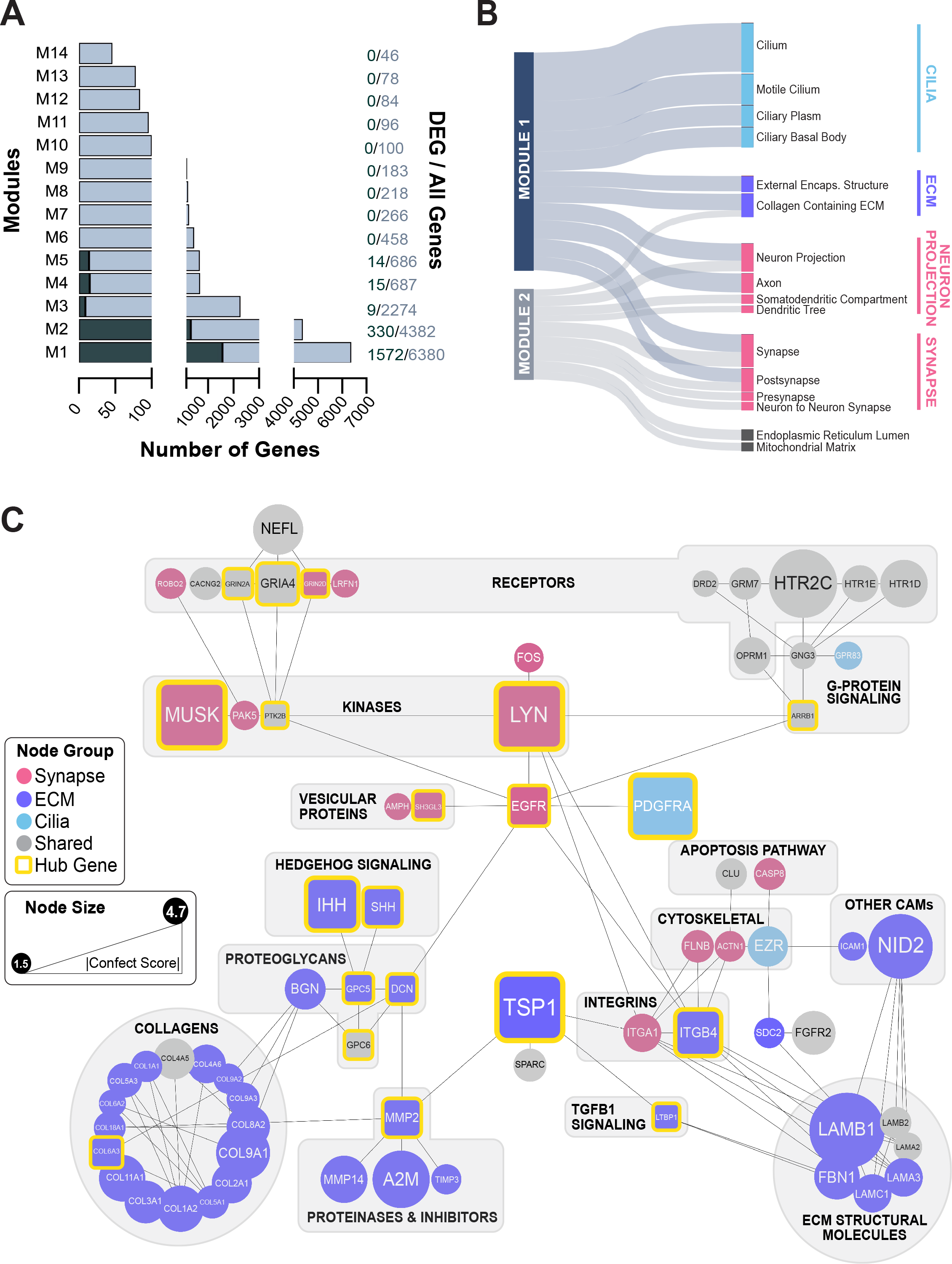
Co-expressed synaptic, ECM, and ciliary encode physically interacting proteins. **(A)** Sizes of all co-expression modules identified among all eBayes-ranked genes following correction for cell line differences. The number of DEGs in each module are shown in green. Module sizes and DEG membership are indicated to the right of each bar. **(B)** Top enriched GO-CC terms associated with each module based on absolute log_10_ p-values, which are represented by the relative thickness of the nodes and edges in the plot. **(C)** Protein-protein interaction network of synapse (pink), ECM (purple), and cilia (blue) associated genes belonging to M1 & M2. ‘Neuronal Projection’ genes were grouped with synaptic genes based on semantic similarity. Nodes in grey are proteins belonging to more than one cellular component category. Node sizes reflect the relative magnitude of absolute confect scores, while edges indicate predicted physical interactions in the brain according to *STRINGDB*. The top hub genes identified by *CytoHubba* using the EcCentricity metric are emphasized in square boxes with yellow borders. Major functional groups are highlighted using labeled grey boxes.

Between the first two co-expression modules, 71 proteins encoded by synaptic, ciliary, and ECM DEGs were predicted to physically interact in the brain based on experimental evidence found in *STRINGDB* (Figure 4C). Topological analysis of this network using the node centrality index EcCentricity in *CytoHubba* identified 20 major hubs spanning each cellular component (Figure 4C; Figure S3). Node eccentricity indicates how easily a protein can be functionally reached by other proteins, indicating it’s centrality of influence in a regulatory network (82). The MCP TSP1 appeared as a major hub based on its magnitude of change, bridging interactions between matrix proteases, structural constituents, and cell-adhesion proteins. Likewise, the epidermal growth factor receptor EGFR was central to several functional ECM and synaptic proteins. The latter included synaptic receptors like the μ-opioid receptor (OPRM1), which appeared alongside dopaminergic, serotonergic, and glutamatergic NMDA and AMPA receptors (GRIN2A, GRIN2D, and GRIA4) that were among the top 20 hubs. EGFR’s relationship with these receptors appeared to be mediated by its association with kinases like LYN, PAK5, and PTK2B and G-protein signaling molecules. EGFR was also shown to interact with the platelet-derived growth factor receptor α (PDGFRA), which is activated in primary cilia and is required broadly during CNS development (114–116).

### A growth factor-MCP regulatory axis is central to synaptic and extra-synaptic changes in gene expression caused by methadone treatment

Since methadone induced changes in this network of co-expressed and interacting synaptic and extra-synaptic DEGs, we explored the regulatory hierarchy of this network further. *IPA* Upstream Regulator Analysis identified TGFβ1 as a principal regulator of the synaptic, ciliary, and ECM DEGs in the first two co-expression modules M1 and M2 (Figure 5A; Data File S3). Upon inclusion into the interaction network described in Figure 4C, TGFβ1was identified as a hub based on its EcCentricity Score (Figure S4; Figure 5B). The top 20 DEGs with the greatest scores comprised a highly interconnected nexus of synaptic, ciliary, and ECM regulatory molecules (Figure 5C). Via interaction with G-proteins and protein tyrosine kinases, OPRM1 linked to a cascade of signaling pathways regulated by the ECM and cilia (e.g., PDGFRA, Hedgehog, and TGFβ1), that were physically linked to both pre-synaptic vesicular trafficking (SH3GL3) and post-synaptic neurotransmitter reception (HTR2C). EGFR and TSP1 were identified as major hub genes by centrality and degree of differential expression. Finally, MCODE cluster analysis of the broader interaction network including TGFβ1 (Figure S4) identified a dense cluster of highly interconnected ECM proteins consisting of this growth factor and the MCPs TSP1, GPC5, GPC6, and SPARC (Figure 5C; Table S4). This finding highlights the centrality of MCPs like TSP1 and growth factors like TGFβ1 in the regulatory response to chronic methadone treatment.

**Figure 5.**
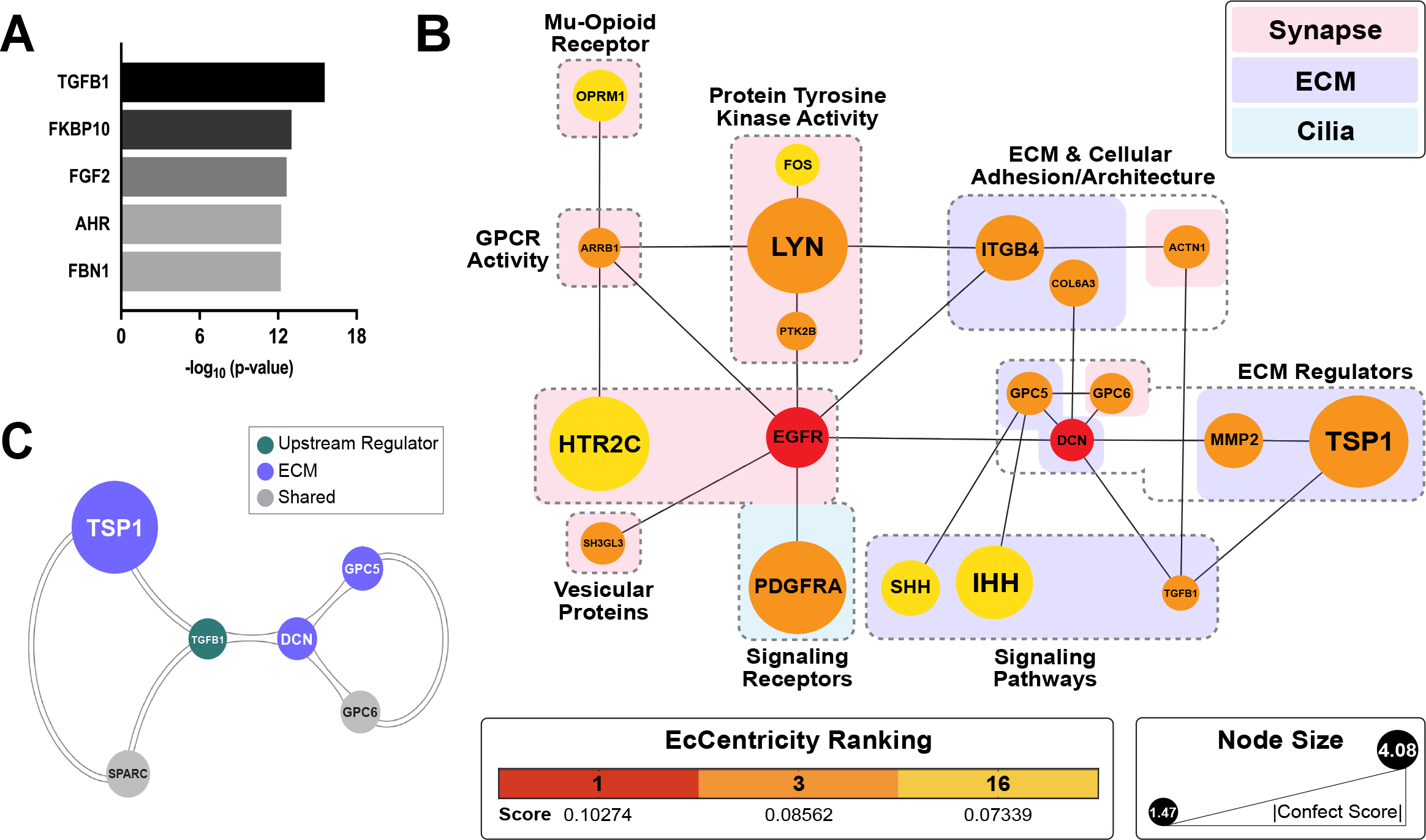
Methadone alters the expression of a central growth factor-MCP regulatory axis. **(A)** Top 5 endogenous protein upstream regulators of the synaptic, ECM, and ciliary genes belonging to modules 1 & 2 in Figure 4B, identified via *IPA*. **(B)** Predicted physical interactions between the top 20 hub genes in a synaptic-extra-synaptic network including the upstream regulator TGFB1. Node colors represent EcCentricity centrality rankings, while node sizes reflect the relative magnitudes of absolute confect scores. Genes are clustered according to their roles in one or more cellular component. **(C)** A highly interconnected network of MCPs and the upstream regulator TGFB1, identified through MCODE (Cluster score = 2.8, node score cutoff = 0.2).

## DISCUSSION

In this study, we investigated the transcriptional effect of chronic methadone treatment on the earliest stages of cortico-genesis modeled in human iPSC-derived hCOs. Our findings provided us with sufficient evidence to posit the following: first, that the previously observed reductions in neural network activity arise from disruptions gene expression programs associated with synapse formation; second, that these synaptogenic changes are brought about by perturbations in developmental signaling pathways originating in the ECM and cilia.

The first of these hypotheses is based on prior evidence of μ-opioid-induced alterations in neural connectivity and communication (20–32). In line with this, we observed significant changes in the expression of genes associated with the pre- and post-synaptic biology of chemical synapses, including the release, reception, or transduction of signals at the synapse (Figure 2). This was congruent with the results of previous multi-electrode array (MEA) and whole-cell patch clamp studies conducted by our lab, which demonstrated dose-dependent attenuations in action potential firing and synaptic transmission (i.e., the frequency and amplitude of spontaneous excitatory post-synaptic currents) in response to methadone in 2 to 4-month-old hCOs (117,118). Taken together with these data, our findings suggest that the methadone-induced suppression of neural network activity arises from perturbations in early pre- and post-synaptic molecular apparatuses that facilitate the establishment of synaptic transmission during prenatal development (87,88). Unexpectedly, given prior evidence, the NES values associated with synaptic gene sets denoted a general trend towards transcriptional upregulation in response to methadone (Figure 1E). Given the indicated physical interactions between synaptic, matrix, and ciliary DEGs, we submit that this upward shift in synaptic gene expression may be a compensatory response to the suppression of essential regulatory mechanisms modulated by the ECM and cilia.

The potential role of the ECM in bringing about changes at the synapse downstream of methadone is reinforced by our finding that the drug alters the expression of a vast matrix regulatory hierarchy (Figure 3A-C, Data File S3). Pivotal to this hierarchy are MCPs, modulators of cell-matrix interactions dynamically expressed at high levels during brain development (106–110). We identified 37 differentially expressed MCPs, spanning several categories of core and matrix associated proteins (Figure 3B-C). Through their known involvement in growth factor signaling, these molecules mediate a balance between ECM structural proteins and the proteases that degrade them, which has downstream effects on cellular adhesion, proliferation, migration, and, ultimately, synapse formation (106–110). Several gene groups belonging to the MCP interactome were also downregulated in response to methadone treatment in our hCOs, including integrins, growth factors and their signaling molecules, collagens, and MMPs (97–104, 122).

Of the canonical differentially expressed MCPs, thrombospondin-1 (TSP1) exhibited the greatest change in expression in response to treatment and appeared as a central hub in a network of co-expressed synaptic and extra-synaptic DEGs (Figure 3C, Figure 4C). TSP1 is a multidomain, multimeric glycoprotein that is secreted into the ECM of synapses. Its domain-specific interactions with growth factors, proteases, and cell surface receptors have proven necessary to proper synaptic architecture and refinement (119–122). A recent study implicated disruptions in the TSP1-TGFβ1-EGFR axis in the loss of synaptic density in rat neuronal cultures following partial μ-opioid receptor activation by morphine (28). Our study identified TGFβ1 as an upstream regulator of the synaptic-extra-synaptic network and both TSP1 and EGFR as a major-hubs in this network based on their degree of change and centrality (Figure 5). This is the first time such an effect has been noted for methadone, a full μ-opioid receptor agonist, in a human-specific model following a treatment regimen clinically relevant for maternal OUD.

Alongside TGFβ1, PDGFRA and SHH/IHH also appeared as major hubs in the synaptic-extra-synaptic regulatory network affected by methadone. PDGFRA was one of the first receptor tyrosine kinases shown to localize to primary cilia, where it mediates signals for directional cell migration and chemotaxis (114–116). In their capacity as sensory projections, primary cilia also contain receptors for Hedgehog, Wnt, Notch, other potent growth factors, integrins, and cadherins (112–114,123). In addition, localization of the soluble SHH and IHH ligands to primary cilia has proven crucial for their signaling, which plays a prominent functional role during synapse formation and circuit assembly (115,116,124). Recent studies have indicated extensive crosstalk between cilia and the ECM, with ciliopathies leading to the dysregulation of ECM proteins such as collagens, laminins, MMPs, and the TGFβ signaling pathway (123,125,126).

Ultimately, we believe that these data contribute to a mechanistic understanding of the neurobehavioral deficits caused by prenatal methadone exposure and provide a foundation upon which to improve pharmacological interventions for OUD in pregnant women.

## ACKNOWLEDGEMENTS

We thank Dr. Kristen Jepsen and the UC San Diego Institute of Genomic Medicine for their assistance with the sequencing conducted for this study.

## Author Contributions

Ila Dwivedi generated the hCO samples, collected the resulting RNA, carried out the processing and systems analysis of the resulting data, and wrote the manuscript. Andrew B. Caldwell provided the requisite base code for the differential expression, gene set enrichment, co-expression network, and protein-protein interaction analyses. Dan Zhou assisted with data pre-processing and conducted the IPA upstream regulator analysis. Both Dan Zhou and Andrew B. Caldwell helped establish the conceptual and mechanistic framework of the paper. Shankar Subramaniam and Gabriel G. Haddad supervised the study and were involved in the overall study design, data interpretation, and revisions of the manuscript.

## Data and Materials Availability

All data required to interpret and evaluate the conclusions presented in this paper are provided in the main figures and supplementary materials. The RNA-sequencing dataset presented in this work is available at the NCBI GEO under the SuperSeries accession GSE210682. Further requests for data and inquiries may be directed to the corresponding author.

