## Supplementary Methods, Figures, and Tables for "Methadone alters transcriptional programs associated with synapse formation in human cortical organoids"

### SUPPLEMENT 1

### SUPPLEMENTARY MATERIALS AND METHODS

#### Human cortical organoid generation

iPSC colonies of identical passage number (across clones and cell lines) were dissociated, collected by centrifugation, and resuspended in mTeSR™1 medium (Stem Cell Technologies, Vancouver, Canada) supplemented with 10 μM SB431542 (Reprocell, Beltsville, MD) and 1μM Dorsomorphin (Tocris, Bristol, UK) to promote neuroepithelial cell identity. Free-floating embryoid bodies were obtained by resuspending iPSCs at a density of approximately 5 x $10^6$ cells per well in a 6-well plate (Corning, Corning, NY) and placing the plate on an orbital shaker at 95 rpm. Y-27632 ROCK Inhibitor (Tocris) was only added for the first 24 hours to promote cell survival during self-aggregation. After three days in this initial suspension, the organoids were shifted to Media1 (M1) for 7 days to initiate neural induction. M1 consisted of a Base Medium (M2) [Neurobasal Medium (Thermo Fisher Scientific, Waltham, MA), 1X GlutaMax, 1X NEAA, 1X Penicillin-Streptomycin (Life Technologies, Carlsbad, CA), 1X Gem21, and 1X N2 (Gemini BioProducts, Sacramento, CA)] supplemented with 10 μM SB431542 and 1 μM Dorsomorphin. Next, the media was changed to Media2-F (M2F) with the addition of 20ng/mL FGF (Stem Cell Technologies) for 7 days, followed by Media2-EF (M2EF) supplemented with 20 ng/mL FGF and 20 ng/mL EGF (PeproTech, Westlake Village, CA) for 6 days. The organoids were then exposed to Media3 for 7 days, which contained growth and neurotrophic factors to encourage neural maturation: 10 μg/mL of BDNF, 10 μg/mL of GDNF, 10 μg/mL of NT-3 (PeproTech), 200 μM L-Ascorbic Acid and 1 mM dibutyryl-cAMP (Sigma-Aldrich, St. Louis, MO). Subsequently, the hCOs were maintained for as long as required in Base Medium M2. All cell culture media was filtered with Tube-Top Vacuum Filters (Corning) to ensure sterility. Growth factors and small molecules were added to the medium the day of media change.

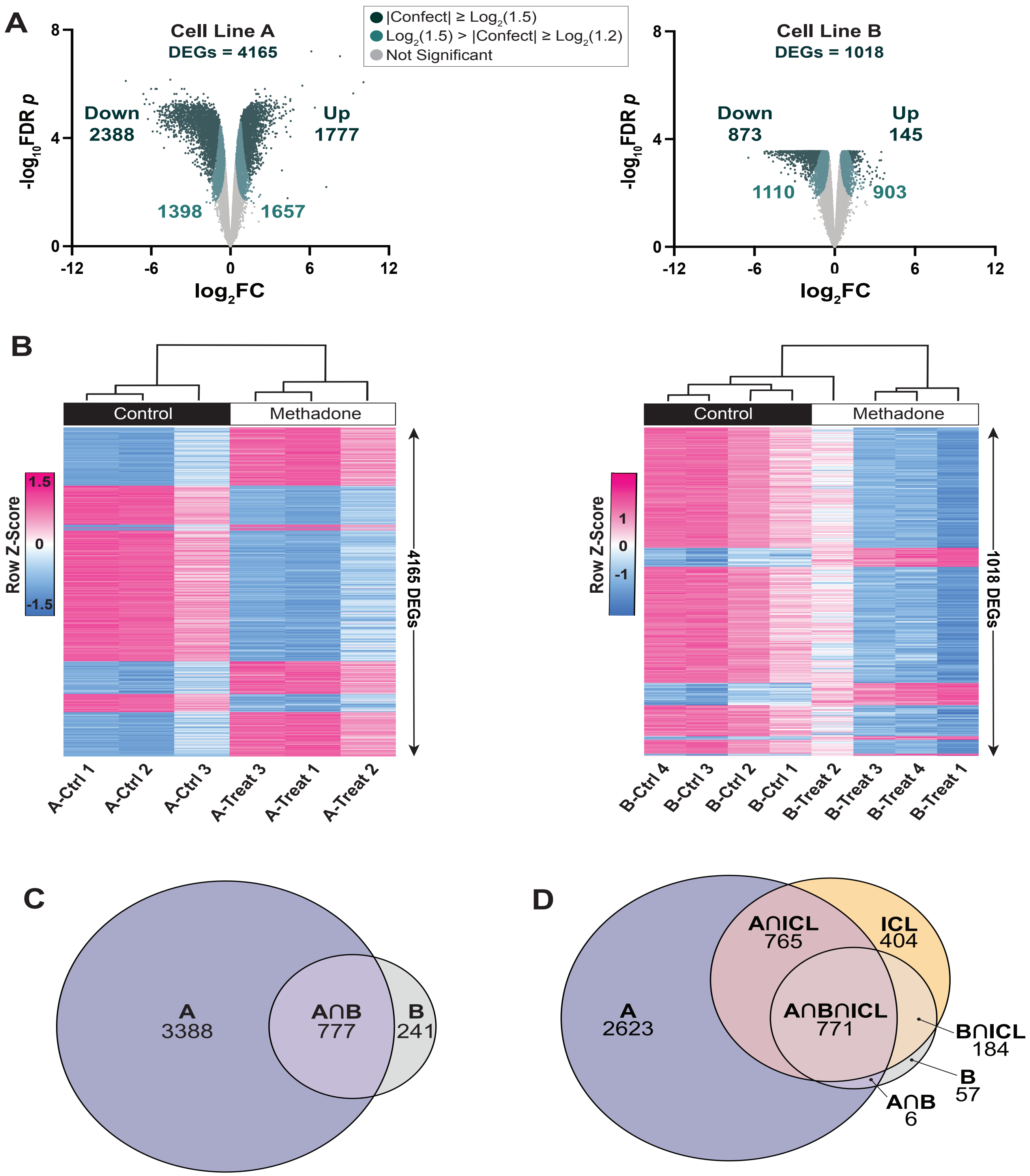


**Figure S1. Robust transcriptional response of individual cell lines to chronic methadone treatment. (A)** RNA-seq volcano plots of expressed genes in cell lines A and B, distinguished by their confident effect size 'confect' score cutoffs. DEGs were genes with |Confect| ≥ $\log_2(1.5)$ and FDR < 0.05, following adjustment for cell-line differences in *limma* (Line A DEGs = 4165, Line B DEGs = 1018). For each gene, $\log_2$ (Fold Change) effect size values and Benjamini-Hochberg adjusted absolute $\log_{10}$ p-values, obtained using the Treat method in *limma*, are shown. **(B)** Heat maps depicting the sample-level expression of DEGs in cell lines A and B, respectively. Expression values are represented as z-scores. **(C)** Overlap of DEGs between cell lines A and B. **(D)** Overlap between DEGs from cell lines A and B, respectively, and DEGs after accounting for cell-line differences inexpression (ICL, independent of cell line).

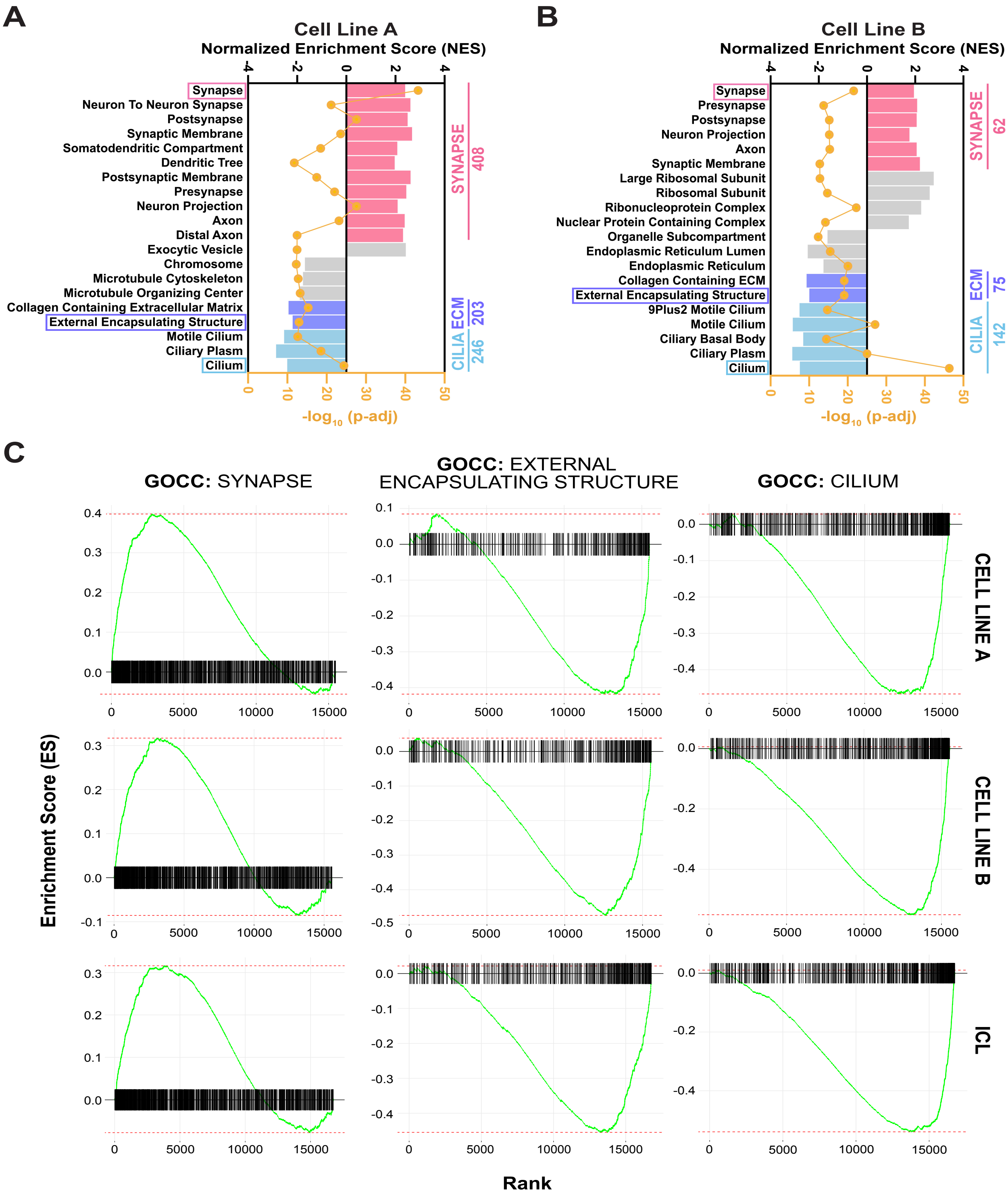


**Figure S2. DEGs in both cell lines are enriched for synaptic, ECM, and ciliary gene expression signatures. (A-B)** Top 20 *fgsea* enriched GOCC gene sets in both cell lines based on absolute $\log_{10}$ FDR-adjusted p values. Gene sets were ranked according to the direction (positive or negative) of their normalized enrichment scores (NES) and grouped by cellular component. Values to the right of each graph indicate the number of DEGs associated with each of the top 3 enriched GO-CC gene sets (synapse, external encapsulating structure (ECM), and cilium). **(C)** *fgsea* enrichment plots for the top 3 enriched GOCC terms in each individual cell line and following adjustment for cell line differences (ICL, independent of cell line). Green lines represent the running enrichment score (ES) for each GO term as the analysis proceeds through the ranked list of expressed genes (black bars) in each dataset, peaking at the final ES score for that term.

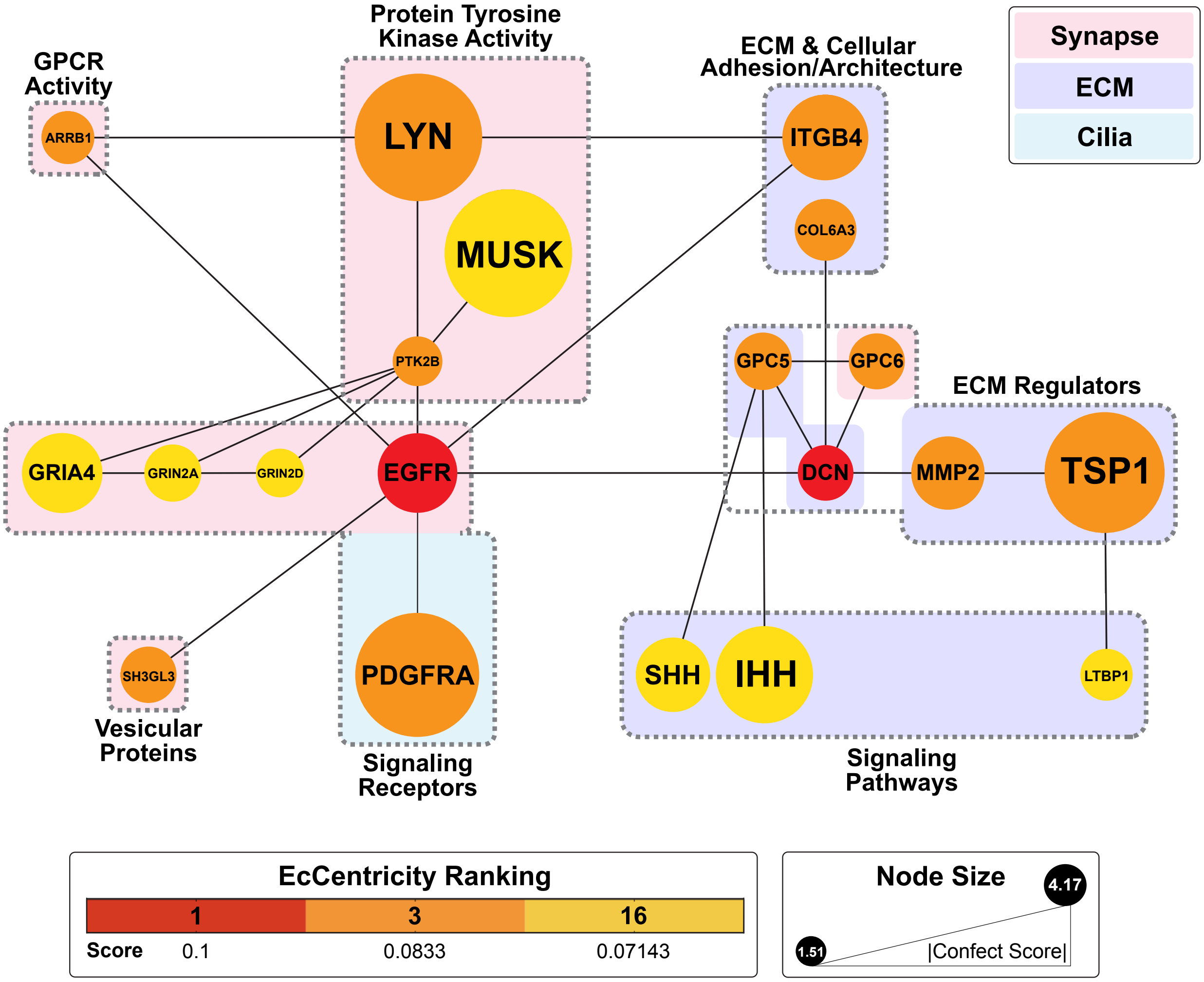


**Figure S3. Proteins encoded by synaptic and extra-synaptic hub DEGs comprise a central regulatory network.** Predicted physical interactions between the top 20 hubs in a network of proteins encoded by co-expressed synaptic and extra-synaptic DEGs. The color of each node represents its centrality ranking according to the EcCentricity scores generated in *CytoHubba.* Both rankings and scores are shown in the legend below the network. Node sizes reflect the relative differential expression of each DEG based on absolute confect scores. Proteins are highlighted in purple, blue, or pink based on their association with synapses, the ECM, or cilia according to the GOCC database, and are clustered into broad functional groups (gray dotted lines).

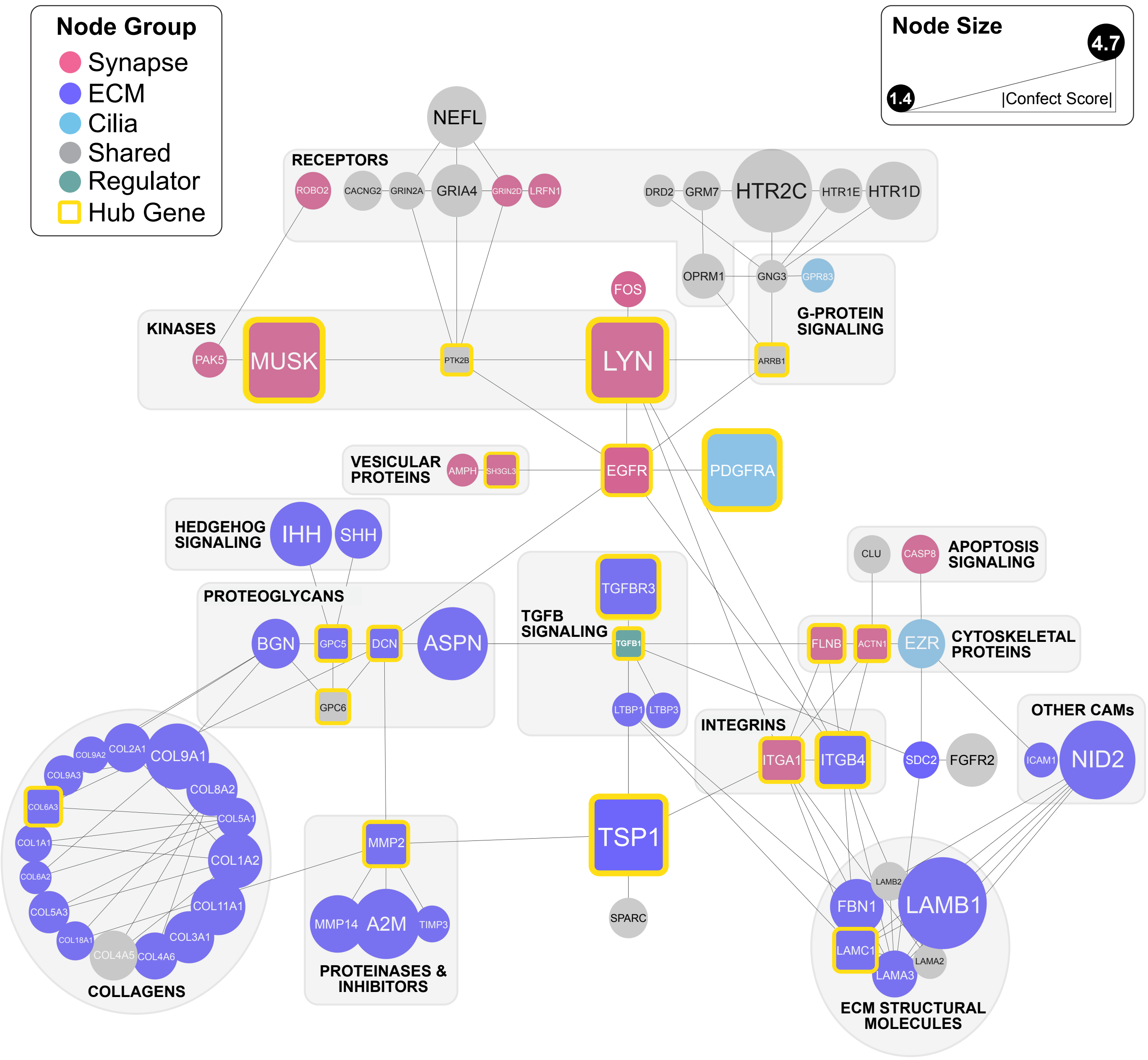


**Figure S4. Synaptic and extra-synaptic protein-protein interaction network including the top upstream regulator TGFB1.** Proteins encoded by the synapse (pink), ECM (purple), and cilia (blue) associated DEGs belonging to the first two co-expression modules are shown, as well as the top upstream regulator TGFB1 (green). Nodes in grey represent genes belonging to more than one cellular component category. Node sizes reflect the relative magnitude of absolute confect scores, while edges indicate predicted physical interactions in the brain according to the STRING database. The top EcCentricity hubs identified by *CytoHubba* are emphasized with square boxes and yellow borders. Major known functional groups are shown in labeled grey boxes.

**Table S1. Quality control information for each sequenced sample.** RNA Integrity Numbers (RIN) and the total number of paired end reads are provided for each sequenced sample. The total reads for each sample are the sum of their forward (read 1) and reverse (read 2) reads.

| | Sample ID | Treatment | RIN | Total Reads[a] |
|---|---|---|---|---|
| **Cell Line A** | A-Ctrl 1 | Control | 9.2 | 106,055,648 |
| | A-Ctrl 2 | Control | 8.4 | 141,760,854 |
| | A-Ctrl 3 | Control | 8.9 | 122,330,614 |
| | A-Treat 1 | Methadone | 8.8 | 97,979,038 |
| | A-Treat 2 | Methadone | 8.2 | 103,402,100 |
| | A-Treat 3 | Methadone | 8.4 | 122,481,702 |
| **Cell Line B** | B-Ctrl 1 | Control | 9.0 | 123,832,834 |
| | B-Ctrl 2 | Control | 9.0 | 103,326,776 |
| | B-Ctrl 3 | Control | 8.1 | 98,315,580 |
| | B-Ctrl 4 | Control | 9.3 | 122,938,118 |
| | B-Treat 1 | Methadone | 8.2 | 95,968,368 |
| | B-Treat 2 | Methadone | 9.0 | 129,797,862 |
| | B-Treat 3 | Methadone | 8.6 | 95,841,676 |
| | B-Treat 4 | Methadone | 7.2 | 138,367,934 |

**Table S2. Quantification of DEGs in each ECM molecular function category.** The number of ECM-associated DEGs belonging to each molecular function category visualized in Figure 3B.

| MatrisomeDB Division | MatrisomeDB Category | Molecule Type | Number of DEGs |
|---|---|---|---|
| Core ECM | ECM Glycoproteins | ECM Assembly Proteins | 5 |
| | | Growth Factor Receptor/Binding Protein | 5 |
| | | Laminin | 5 |
| | | Other Cell Adhesion Molecules | 4 |
| | | CCN Family Proteins | 3 |
| | | Fibulin | 3 |
| | | Integrin Signaling/Signal Transduction | 2 |
| | | Nervous System Enriched ECM Proteins | 2 |
| | | Collagen-Like/ Related | 1 |
| | | Fibrillin | 1 |
| | | Integrin | 1 |
| | | Nidogen | 1 |
| | | Protease Regulation | 1 |
| | | Thrombospondin | 1 |
| | Collagens | Fibril Forming | 7 |
| | | Fibril Associated | 5 |
| | | Network-Forming | 3 |
| | | Beaded Filament | 2 |
| | Proteoglycans | Extracellular Proteoglycan/SLRP | 6 |
| | | Cell-Surface Proteoglycans | 3 |
| | | Hyaluronan & Proteoglycan Link Protein | 1 |
| | | Basement Membrane Proteoglycan | 1 |
| Matrisome Associated | ECM Regulators | Protease/Proteinase | 16 |
| | | ECM Crosslinking/Remodeling Enzymes | 8 |
| | | Protease/Proteinase Inhibitor | 7 |
| | ECM Affiliated Proteins | Annexins | 6 |
| | | Nervous System Enriched ECM Proteins | 4 |
| | | Other Basement Membrane Components | 3 |
| | | Immune/Inflammatory Response | 3 |
| | | Growth Factor Receptor/Binding Protein | 2 |
| | | Growth Factor | 1 |
| | | BMP Signaling | 1 |
| | | Galectin | 1 |
| | Secreted Factors | WNT Signaling | 4 |
| | | Growth Factor Signaling | 2 |
| | | Angiopoietin Signaling | 2 |
| | | BMP Signaling | 2 |
| | | Hedgehog Signaling | 2 |
| | | Annexin Signaling | 1 |
| | | Apoptosis Signaling | 1 |

**Table S3. Top 10 enriched GOCC categories for co-expression modules 1 and 2.** *fGSEA* using the GOCC database yielded the enrichment of synaptic, ciliary, and ECM associated transcriptional signatures among all genes in the top 2 co-expression modules. This data was used to generate the Sankey plot in Figure 4B.

| | Pathway | $-log_{10}$ (p-adj) | NES |
|---|---|---|---|
| **Module 1** | Cilium | 34.185 | -3.279 |
| | Motile Cilium | 21.080 | -3.391 |
| | Ciliary Plasm | 14.217 | -3.129 |
| | Ciliary Basal Body | 13.855 | -3.073 |
| | Axon | 13.370 | 2.654 |
| | Synapse | 12.314 | 2.212 |
| | Neuron Projection | 12.179 | 2.210 |
| | Collagen Containing Extracellular Matrix | 11.417 | -2.666 |
| | External Encapsulating Structure | 10.678 | -2.471 |
| | Postsynapse | 9.578 | 2.463 |
| **Module 2** | Synapse | 10.293 | 2.253 |
| | Neuron Projection | 7.349 | 2.079 |
| | Mitochondrial Matrix | 7.349 | -2.583 |
| | Postsynapse | 6.146 | 2.227 |
| | Somatodendritic Compartment | 6.146 | 2.152 |
| | Neuron To Neuron Synapse | 5.668 | 2.335 |
| | Endoplasmic Reticulum Lumen | 5.534 | -2.439 |
| | Presynapse | 5.509 | 2.186 |
| | Collagen Containing Extracellular Matrix | 4.578 | -2.278 |
| | Dendritic Tree | 4.470 | 2.065 |

**Table S4. Networks of highly interconnected nodes associated with ECM structure and regulation.** Highly interconnected sets of nodes identified in the synaptic-extra-synaptic network including the upstream regulator TGFB1 (Figure S4) using *MCODE*. For each cluster, the number of nodes and edges are provided, including a score calculated by multiplying the density of the cluster by the number of nodes.

| Cluster | Score | Nodes | Edges | Node IDs |
|---|---|---|---|---|
| **1** | 4.50 | 5 | 18 | COL9A3, COL2A1, COL9A1, BGN, COL9A2 |
| **2** | 3.33 | 4 | 10 | COL5A3, COL1A2, COL5A1, COL1A1 |
| **3** | 2.80 | 6 | 14 | TGFB1, DCN, SPARC, GPC6, TSP1, GPC5 |

**Data File S1. *REVIGO* tree maps generated for DEGs as input for *CirGO* visualization.** Each sheet contains a list of the enriched non-redundant GO-MF terms associated with the synaptic, ECM, and ciliary DEGs, respectively. Terms without a representative "parent" were assigned one based on the highest significant term in the GO hierarchy. This data was then used to generate the *CirGO* plots seen in Figures 2 and 3.

**Data File S2. Molecular Functions of ECM and Synaptic DEGs.** Categorization of the DEGs associated with the synapse (n = 166) and the ECM (n = 129). *Matrisome DB* was used to inform ECM DEG identities. Both synaptic and ECM genes were grouped according to information available in the *OMIM, GeneCards,* and *NCBI Gene* databases**.** File also includes MCPs identified through published reviews. MCPs were identified as classically found in the brain (Classical), included through expansions of the MCP definition (Expanded), or those that have been proposed but are still under debate (Debated).

**Data File S3. *IPA* Upstream Regulator Analysis output.** Unfiltered output of all upstream regulators identified by Qiagen *IPA* for synaptic, ECM, and ciliary DEGs in co-expression Modules 1 and 2. Filtered list excluding exogenous toxins, chemicals, and drugs is also included.
